# Tonic interferons defend against respiratory viruses in primary human lung organoid-derived air-liquid interface cultures

**DOI:** 10.64898/2025.12.10.693424

**Authors:** Rinu Sivarajan, Paul C Kirchgatterer, Jan Lawrenz, Eszter Tanner-Matiz, Jessica Lindenmayer, Verena Renz, Tapan Joshi, Heike Oberwinkler, Thorsten Walles, Giorgio Fois, Alexander Kleger, Manfred Frick, Jan Münch, Moritz M Gaidt, Maria Steinke, Konstantin MJ Sparrer

**Affiliations:** Institute of Molecular Virology, Ulm University Medical Center, Ulm, Germany; Research Institute of Molecular Pathology, Vienna BioCenter, Vienna, Austria; Core Facility Organoids, Ulm University Medical Center, Ulm, Germany; Department of Oto-Rhino-Laryngology, Head and Neck Surgery, University Hospital Würzburg, Würzburg, Germany; Department of Thoracic Surgery, University Medicine Magdeburg, Magdeburg, Germany; Institute for General Physiology, University of Ulm, Ulm, Germany; Institute of Molecular Oncology and Stem Cell Biology, Ulm University Hospital, Ulm, Germany; German Center for Neurodegenerative Diseases (DZNE), Ulm, Germany

## Abstract

Innate defences of the respiratory epithelium are the first barrier against incoming respiratory viruses. To understand the contribution of both basal (tonic) and induced interferon (IFN) to antiviral defences in a physiologically relevant system, we established air-liquid interface (ALI) cultures of primary human bronchial epithelium (HBE) and small airway epithelium (HSE). Via an organoid intermediate stage, the limited healthy donor material was expanded while preserving stemness and subsequently differentiated. Characterisation by spatial and transcriptomic analyses showed that the cellular diversity and architecture of our ALI cultures were comparable to native human lung epithelium. Upon infection with relevant human respiratory pathogens, such as Human Rhinovirus (HRV16) and human Coronaviruses (229E and NL63), only HRV16 induced a strong and early type I and III IFN response, leading to its eventual clearance from the cultures. Depletion of tonic type I/III IFNs using neutralising antibodies or scavengers reduced expression of levels of IFN-stimulated genes and increased infectious HRV production by ∼7-10-fold. Taken together, we present a method for generating primary lung epithelial cultures that retain their IFN status, demonstrate clearance of HRV by innate defences, and highlight the importance of tonic IFN in early antiviral defences.

**IMPORTANCE:** Mild respiratory viral infections, for example, with human common cold coronaviruses or rhinoviruses, are a massive cause of human morbidity. The respiratory tract is the primary entry route for these viruses and also the contact site for initial innate immune defences. Here, we show that primary human lung epithelial cell-derived air-liquid interface cultures mimic the architecture and cell composition of native human lung epithelium, and retain both induced and tonic interferon (IFN) responses. Notably, our data show that the model’s innate immune defences are sufficient to clear human Rhinovirus (HRV) infections, which are characterised by rapid and robust IFN responses. Finally, depletion of tonic IFNs led to a marked increase in HRV infection. Thus, our research suggests that tonic low levels of IFNs contribute to the epithelial defence against viruses, maintaining the tissue’s immune readiness. Failure to maintain these tonic IFN levels increases the susceptibility towards infections.

## INTRODUCTION

Infections of the respiratory tract are a significant burden to human health, and consistently ranked among the top 10 leading causes of death worldwide over the last few decades (1). Endemic respiratory viruses such as the human Rhinoviruses (HRVs) and the human coronaviruses (HCoV-229E and HCoV-NL63) are major contributors to seasonal colds. Together, these viruses represent the most common cause of respiratory infections, making them among the most frequent viral encounters (2). While they mainly cause mild infection in healthy adults; children, immunocompromised, and elderly individuals may develop severe course of the disease (3–6). The entry site for these respiratory pathogens is the airway epithelium, a thin continuous layer of specialised epithelial cells that line the respiratory tract. The majority of HRV strains infect cells via the intercellular adhesion molecule-1 (ICAM-1); a small group also uses the low-density lipoprotein receptor (LDLR) or the Cadherin-like family member 3 (CDHR3) (7–10). Common cold coronaviruses infect cells via angiotensin-converting enzyme 2 (ACE2) (e.g. hCoV-NL63) or aminopeptidase N (hCoV-229E) as entry receptors that are expressed throughout the airway epithelium (11, 12).

Besides the upper conducting airways, the lower respiratory epithelium serves as the first barrier of immune defence against invading viruses, ideally enabling their rapid clearance with minimal impact on the host. (13). It is mainly composed of ciliated cells, secretory cells, goblet cells, basal cells, club cells and minor populations of pulmonary neuroendocrine cells, pulmonary ionocytes, hillock cells and tuft cells (14). Ciliated cells are the most abundant and are responsible for the movement of mucus and trapped particles via coordinated ciliary motion. Secretory cells and goblet cells, interspersed among ciliated cells, produce mucus, which forms a protective layer that traps dust, pathogens, and other particulate matter. While the mucus layer serves as a physical barrier, the ciliary motion promotes mechanical clearance of mucus and trapped particles from the respiratory tract. Basal cells serve as stem cell progenitors, capable of differentiating into other epithelial cell types to maintain tissue integrity and regeneration (14–16). Additionally, epithelial cells secrete antimicrobial peptides, such as defensins and lysozyme. The epithelial cells also express pattern recognition receptors (PRRs) that detect pathogen-associated molecular patterns (PAMPs), leading to the release of pro-inflammatory cytokines, such as Interferons (IFNs). Secretion of IFNs induces the expression of IFN-stimulated genes (ISGs), many of them with well-known anti-viral functions, setting the cells in an anti-viral state (17–19). Of note, whereas Type I and II IFNs (e.g. IFN-α, IFN-β, IFN-γ) are associated with systemic anti-viral responses, Type III IFNs (IFN-λ) was proposed to have a more specialised role in protecting mucosal surfaces (20–23).

Recent studies have highlighted tonic or basal IFN as a means to maintain ‘immune readiness’ in the absence of overt infection, shaping the basal innate immune defences and inflammatory state of the airway epithelium (24). These tonic interferons, primarily type I IFNs such as IFN-α and IFN-β, maintain a low-level transcriptional program that primes cells for rapid antiviral responses while also contributing to immune homeostasis (25–28). Decreased levels of tonic IFNs have been associated with increased susceptibility to respiratory viruses such as severe acute respiratory syndrome coronavirus 2 (SARS-CoV-2), and dysregulation of tonic IFNs has been implicated in autoimmune diseases, chronic inflammation, and tumour immune evasion (29–31). Despite their importance for defence, the impact of tonic IFNs on anti-viral defences at the airway epithelium remains poorly understood. A major limitation is that most studies rely on immortalised cell lines or 2D-cultured primary bronchial epithelial cells, which do not accurately recapitulate the defence responses of the human airway epithelium. In recent years, human primary airway epithelial cells cultured at the air–liquid interface (ALI) have emerged as a powerful tools for studying respiratory host-pathogen interaction (32–34). Even though some challenges, such as donor availability, variability, and a short lifespan come along with these models, they closely mimic the physiology of the human airways, providing complex and human-focused in vitro models for development of novel therapeutic approaches (35–37). To overcome these bottlenecks with existing techniques we established here a protocol to generate ALI cultures from normal, commercially available human primary lung epithelial cells (37, 38). To overcome the limitation of cell availability and donor availability, we generated lung organoids that retain stemness and differentiated them at ALI making them similar to native human bronchial epithelium regarding cell composition and architecture. Using this model, we here show that HRV, which induces the highest level of initial IFNs, is cleared from the cultures after ∼4 weeks. Furthermore, depleting of tonic type I and type III IFN reduces basal ISG levels and weakened epithelial defences against HRV by more than 10-fold. Thus, our data show that tonic IFNs contributes to the defences of the respiratory epithelium.

## RESULTS

### Establishing an organoids-intermediate primary ALI culture lung model

Current primary cell-derived ALI cultures of lung epithelium are limited by the availability of donor material, and cells beyond passage three rarely differentiate (39). To address this limitation, we developed a method to amplify primary cells while retaining their stemness via an organoid intermediate state (37). Cells derived from the organoids could either be cryopreserved for long-term use, passaged up to 12 times or directly seeded onto transwell inserts for ALI culture generation (Fig. 1A and S1A). In brief, commercially available primary normal human bronchial epithelial cells or small human airway epithelial cells were grown from an early passage with BME type II and organoid growth medium. On the fifth day, the cells displayed an organoid-like, rounded morphology and stained positive for Cytokeratin 5 (CK5), identifying them as basal cells (Fig. 1B and C). To assess the potential of these continuously passaged basal cells to differentiate into ALI cultures, the organoids were dissociated into single cells and seeded onto transwell inserts, allowing them to expand and form a monolayer (Fig. 1A). Afterwards, the apical medium was removed to create an air-liquid-interface.

**Figure 1.**
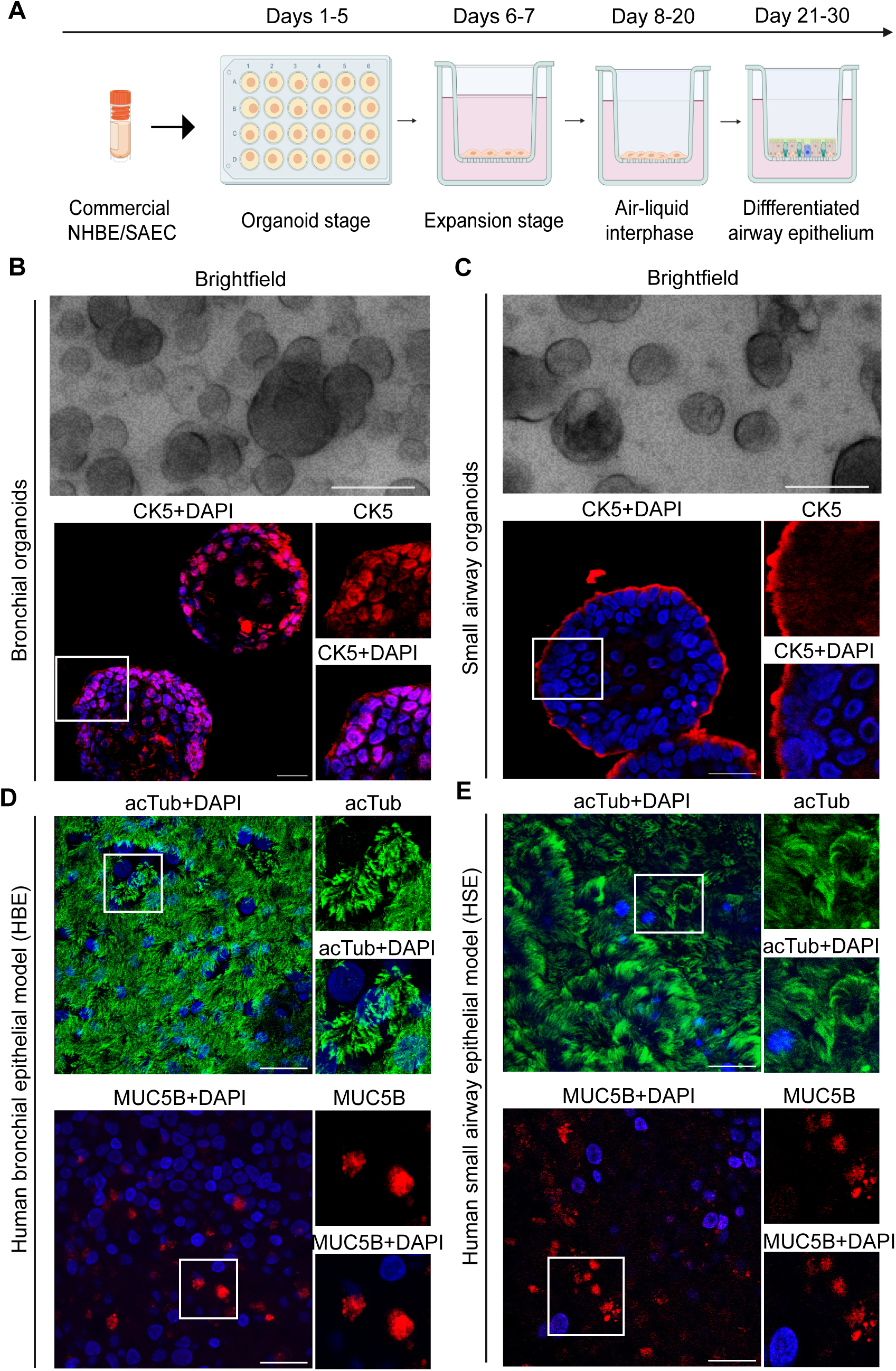
Generation and characterisation of human airway epithelial models from primary human lung epithelial cells. (**A**) Schematic representation of the workflow of human airway epithelial model generation via organoid intermediate using human primary lung epithelial cells. (**B**-**C**) Representative immunofluorescence and brightfield images of bronchial and small airway organoids in brightfield microscopy or confocal scanning microscopy of formalin fixed paraffin embedded (FFPE) 5 µm sections stained for basal cells (CK5+, red). Nuclei, DAPI (blue). Scale bars, 100 µm (bright field microscopy) and 25 µm (confocal microscopy). (**D**-**E**) Representative laser scanning confocal microscopy images of differentiated human bronchial and small airway epithelial models stained for ciliated cells (acTub+, green) and secretory cells (MUC5B+, red). Nuclei, DAPI (blue). Scale bars, 25 µm.

The cultures derived from NHBE and SAEC organoids differentiated into a human bronchial epithelium (HBE) or human small airway epithelium (HSE), respectively. Whole mount staining of acetylated tubulin (acTub) shows kinocilia present on the ciliated cells across the apical surface, and MUC5B staining indicated the presence of secretory cells (Fig. 1D and E). Trans-Epithelial Electrical Resistance (TEER) measurements from ALI cultures generated from low and highly passaged organoids confirm their capacity to form an intact barrier. The TEER values ranged from 500 to 900 Ω · cm2, which is within physiological levels (Fig. S1B). Lung organoids up to passage 12 were tested still showed mucociliary phenotype, suggesting that they differentiated (S1A). Live-cell imaging demonstrated the presence of functional cilia, characterised by the typical synchronised movement of Dynabeads Protein G particles (Video S1).

Taken together, this data demonstrates that ALI cultures were successfully differentiated from human primary bronchial epithelial cells or small airway epithelial cells via organoid-intermediate state, which allows easy amplification of the source material without diminishing its differentiation potential.

### Organoid-intermediate ALI cultures resemble the architecture and composition of native human lung epithelium

The respiratory epithelium is a pseudostratified columnar epithelium, with the columnar ciliated cells framed on the apical side, interspaced by mucus-producing goblet and secretory cells, with a final basolateral layer formed by basal cells (32). Z-axis sections of formalin-fixed paraffin-embedded (FFPE) of HBE ALI cultures confirmed the presence of kinocilia present on the ciliated cells (acTub) on the apical surface. The major gel-forming mucins are MUC5AC and MUC5B secreted by goblet (MUC5AC) and secretory cells (MUC5B) (40). In HBE, the MUC5AC+ and MUC5B+cells cluster towards the apical side (Fig. 2A). Finally, CK5+ basal cells were exclusively found on the basal side of the epithelium (Fig. 2A). A similar polarised epithelial architecture was evident for the HSE ALI cultures, where ciliated cells, goblet, secretory and basal cells were stained using their respective cell markers (Fig. 2B). To determine how close our models resemble native human lung epithelium, we compared the presence and spatial organisation of cellular markers detected in ALI cultures to native human bronchial tissue sections. Similar to the ALI cultures, the ex vivo tissue displays acTuB positive cells ciliated cells, MUC5AC positive goblet cells, MUC5B expressing secretory cells, and CK5-stained basal cells (Fig. 2C).

**Figure 2.**
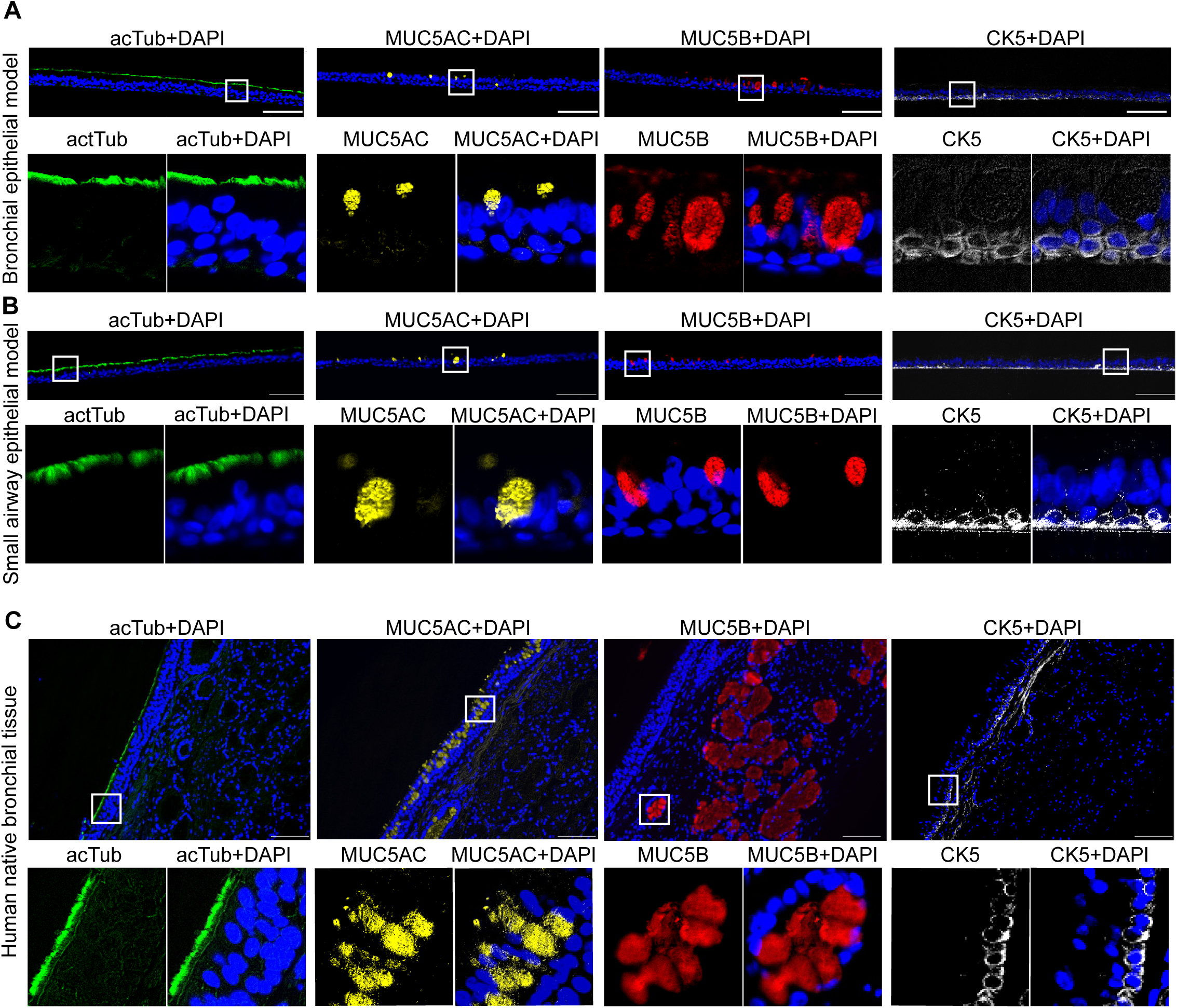
Cellular marker comparison of human airway epithelial models to native human bronchial tissue. (**A**) Representative laser scanning confocal microscopy images of 5 µm FFPE z-sections of the human bronchial epithelial model. Stained for ciliated cells (acTub+, green), goblet cells (MUC5AC+, yellow), secretory cells (MUC5B+, red) and basal cells (CK5+, grey). Nuclei, DAPI (blue). Scale bars, 100 µm. (**B**) Representative laser scanning confocal microscopy images of 5 µm FFPE sections of human small airway epithelial models stained with acTub+ (green), MUC5AC+ (yellow), MUC5B+ (red), and CK5+ (grey). Nuclei, DAPI (blue). Scale bars, 100 µm. (**C**) Representative laser scanning confocal microscopy images of 5 µm sections of native human bronchial tissue stained with acTub (green), MUC5AC (yellow), MUC5B (red), and CK5 (grey). Nuclei, DAPI (blue). Scale bars, 100 µm.

This data indicates that the primary architecture of both the HBE and the HSE models is similar to that of the native human lung.

### HSE and HBE models express respiratory epithelium-associated genes

Different cell types of the human airway epithelium are characterized by specific gene expression patterns (41). Expression of *FOXJ1* is characteristic of ciliated cells, whereas *KRT5* expression marks basal cells. *MUC5AC* and *MUC5B* are expressed by goblet and secretory cells, respectively (42). Both the HBE and HSE models show marker gene expression assessed by qPCR, indicating successful differentiation (Fig. 3A). To comprehensively evaluate the expression patterns of cell-type-specific and respiratory epithelium-associated genes in the HSE and HBE models, we mapped bulk RNAseq across three individual inserts to the healthy lung atlas (Data S1) (14, 41). Cibersort assisted analyses indicated the presence of transcriptional patterns from various cell types, including ciliated, basal, goblet, and secretory cells (Fig. 3B, Fig. S2A, Data S2). The majority of cells were ciliated, followed by secretory and basal cells. Additional cell types, such as serous cells found in submucosal glands, and intermediate cell populations, such as deuterosomal cells, reported as precursors of ciliated cells, which are otherwise difficult to distinguish using immunofluorescence, were also detected using RNAseq. By selectively clustering ciliated, secretory, serous, deuterosomal and basal cell types the ratios of the major cell types in the models were determined (Fig. 3B, Data S3) indicating that overall cell composition was similar between both models. However, principal component analysis of the data suggested that the gene expression profile still differs significantly between HBE and HSE models (Fig. 3C). This is also illustrated by the Top 40 genes by variance (Fig. 3D, Data S4). Gene ontology analyses revealed that genes that are highly expressed in the HSE model are largely part of the mitochondrial respiratory chain, whereas those associated with the HBE model are innate immune and anti-microbial defence genes (Fig. S2B, Data S5). In line with visualisation of basal-level expression of ISGs, mucosal defence, and immunity-associated genes, the HBE model shows higher expression of several prominent defence genes, including defensins, well-known anti-viral ISGs (TRIM5, tetherin/BST-2), and mucins (Fig. 3E-G, Data S5).

**Figure 3.**
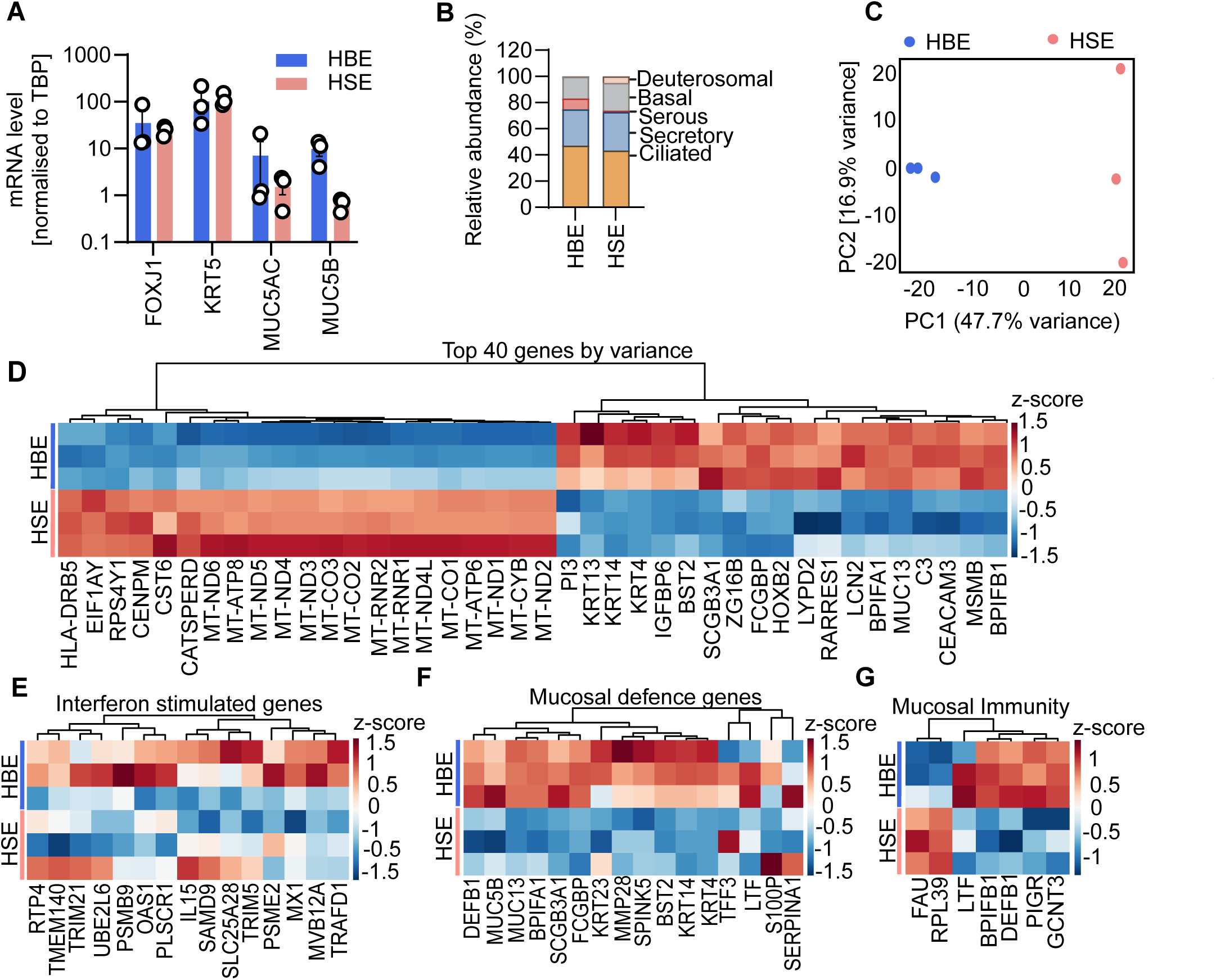
Characterisation of human airway epithelial models at RNA-level. (**A**) mRNA levels of major epithelial marker RNAs, as indicated, assessed by qPCR. Ciliated cells (FOXJ1), basal cells (CK5), goblet cells (MUC5AC) and secretory cells (MUC5B). Normalised to housekeeping gene TBP. The bars represent the mean of n=3±SEM (independent experiments). (**B**) Transcriptomic data of human airway models mapped to the human healthy airway atlas to identify cell types. The data shows the relative abundance of individual cell types from three independent human bronchial and small airway models aggregated from the CIBERSORTx output (Data 2). (**C**) Principal Component Analysis (PCA) of the data in (B). (**D**) Heatmap representation (coloured by z-score) of the top 40 genes by variance showing distinct gene expression patterns between the human bronchial and small airway epithelial models. (**E-G**) Heatmap representation (coloured by z-score) of representative basal type I IFN stimulated genes (E), mucosal defence genes (F) or mucosal immunity Genes (G) in the expression cluster in the human airway models. Levels of the individual biological triplicates are shown.

Taken together, the transcriptome analyses confirm successful differentiation of both models and indicate that the HBE exhibited elevated expression of defence factors, suggesting a more activated state than the small airway epithelium.

### Human bronchial epithelial models self-clear HRV16 in long-term cultures

To analyse the interplay between respiratory virus infections and the innate immune system of the lung epithelium, we infected the HBE model with three clinically prevalent respiratory viruses: human Rhinovirus A (HRV16) and the human coronaviruses 229E and NL63. The HBE model was chosen because it represents the entry tissue for these three viruses. Quantification of viral RNA in apical supernatants showed that all three viruses productively infected the models (Fig. 4A-C). Next, HRV16-infected cultures were treated with Rupintrivir (Rup 3 µM), and 229E- and NL63-infected cultures were treated with Remdesivir (Rem 10 µM) or Camostat Mesylate (CM 20 µM) starting 4 hours post-infection. Rupintrivir significantly reduced HRV16 RNA levels by ∼200-fold at day 4 (Fig. S3A). Both Remdesivir and Camostat Mesylate inhibited 229E by ∼ 20-fold and NL63 by ∼300-fold (Fig. S3B-C). Of note, HRV16 RNA in the supernatant peaked around day 4 and declined thereafter until 16 dpi (Fig. 4A). In contrast, 229E and NL63 RNA remained at high levels up to day 16 post-infection, suggesting that an equilibrium was reached resembling a quasi-persistent infection (Fig. 2B and C). None of the infections were accompanied by large-scale cell death or visible ciliary cessation (Video S2-5). Since cytotoxicity was absent, we further investigated the decline in HRV16. Long-term analyses of HRV16 replication in the HBE model revealed that the RT-qPCR cycle levels of the viral RNA in the supernatant (SN) reached background levels within 14-35 days, depending on the donor, indicating clearance (Fig. 4D). To verify that the infected HBE cultures fully recovered and did not release infectious virus we inoculated H1-Hela cells with the apical SN from human bronchial epithelial models after 31 days of HRV16 infection. The H1-Hela cells showed no cytopathic effect after 3 days, indicating the absence of infectious virus particles. As a positive control we used HRV16 infected cells (Fig. S4A). This clearance was accompanied by pronounced MUC5B secretion; however, the ciliated cell layer (acTub) remained intact (Fig. S4B). Overall, the integrity of the models was not compromised, as shown by the FITC-dextran membrane permeability assay and TEER measurements at 35 days post-infection (Fig. S4C-D). This data indicates that the HBE models can clear a Rhinovirus infection.

**Figure 4.**
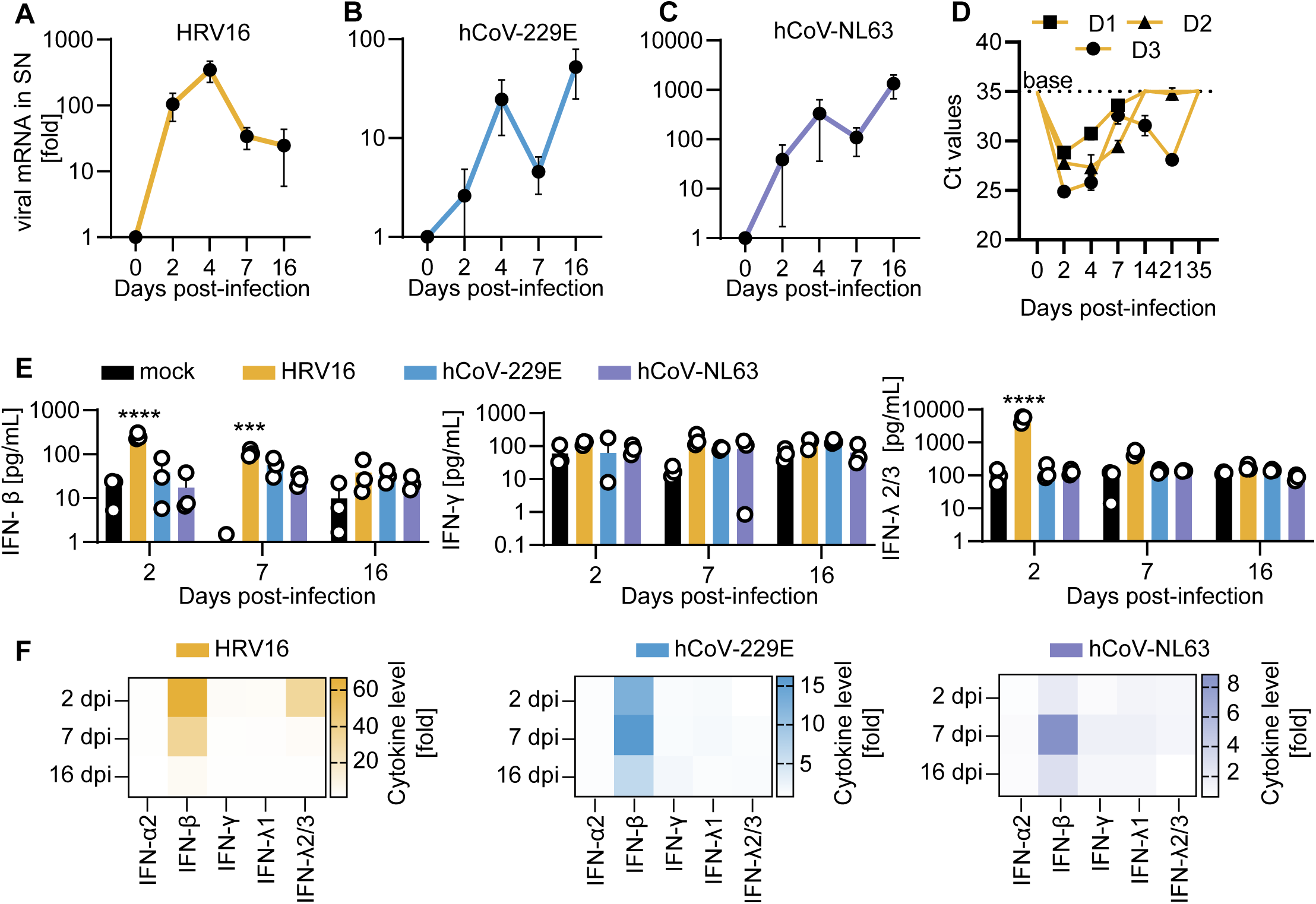
Human bronchial epithelial models support common cold respiratory viral replication and enable long-term monitoring of infection kinetics. (**A-C**) Quantification of the viral RNA levels by qPCR of HBE models infected from the apical side (MOI 0.01, indicated viruses) for 3 h. At indicated days post infection, apical washes were collected. Dots represent the mean of n=3±SEM (independent experiments). (**D**) Quantification of the viral RNA levels by qPCR of HBE models infected with HRV16 (MOI 0.01). At indicated days post infection, apical washes were collected. Shown are the raw Ct values obtained. The baseline (base) is at the Ct levels in the 0 control. Dots represent the mean n=3±SEM (donors). N.d., below detection limit (grey line). (**E**) Absolute cytokine levels as assessed by multiplex ELISA (Legendplex) of the models in (A). Bars represent the mean of n=3±SEM (independent experiments). Two-way ANOVA with Tukey’s multiple comparisons test. ****p< 0.0001, *** p= 0.0003. (**F**) Heatmap depiction of the data in (E) separated by virus and calculated as fold. The data are shown as mean of three independent experiments.

### Rhinovirus activates strong and early cytokine defences in HBE

Since viral clearance in this model depends solely on epithelial innate immune responses, we analysed temporal differences in cytokine release using multiplex cytokine ELISA. Only HRV16, but not the two Coronaviruses, induced a strong and significant cytokine response with higher levels of IFN-α, IFN-β, IFN-λ2/3, IFN-λ1, IL-8, IFN-γ, IL-1β, IL-6, TNF-α and IP-10 (Fig. 4E and Fig. S5). The cytokine responses against HRV16 peaked on day 2 and then declined. In contrast, the hCoV infections induced not only a weaker but also a more delayed response, peaking at day 7 post-infection (Fig. 4F).

Taken together, HRV infection is accompanied by strong and early cytokine response, whereas the innate immune response against the human coronaviruses was much more subdued. This suggests that an early and strong cytokine response mediates HRV clearance.

### Tonic interferons in lung epithelium serves as a major barrier against HRV16

Our data showed that the HBE model may be in a pre-activated state, with prominent IFN stimulated genes highly expressed at basal states (Fig. 3). Indeed, in apical washes various IFNs were readily detectable. On average the HBE model secreted ∼28 pg/mL of IFN-α2, ∼10 pg/mL of IFN-β, ∼18 pg/mL of IFN-γ, ∼150 pg/mL of IFN-λ1, and ∼120 pg/mL of IFN-λ2/3 (Fig. 5A). To determine the impact of the tonic IFN on HRV16, we depleted them by pre-treating the HBEs with neutralising antibodies against IFN-β, IFN-λ antibody cocktail or B18R, a Type I IFN scavenger for 2 days prior to the infection (43). Afterwards the treatments were removed and the cultures infected (Fig. 5B). Depletion of the tonic IFN was successful as shown by ∼31% reduced basal MX1 levels in both B18R, and anti-IFN λ treated HBE cultures and ∼10% reduction in MX1 levels in anti-IFN-β (Fig. 5C). Together the anti-IFN-β, and anti-IFN-λ treatment resulted in ∼2-fold higher intracellular viral RNA and B18R treatment resulted in ∼ 5 fold higher HRV16 RNA content compared to untreated controls (Fig. S6). Importantly, infectious virus yields in the apical supernatants assessed after 24 h post-infection increased markedly in IFN-depleted cultures (Fig 5D). On average, depletion of type I IFN by neutralising antibody, depletion of type III or scavenging of IFN by B18R, increased infectious virus yields by ∼3, 10 and 7-fold respectively (Fig. 5D).

**Figure 5.**
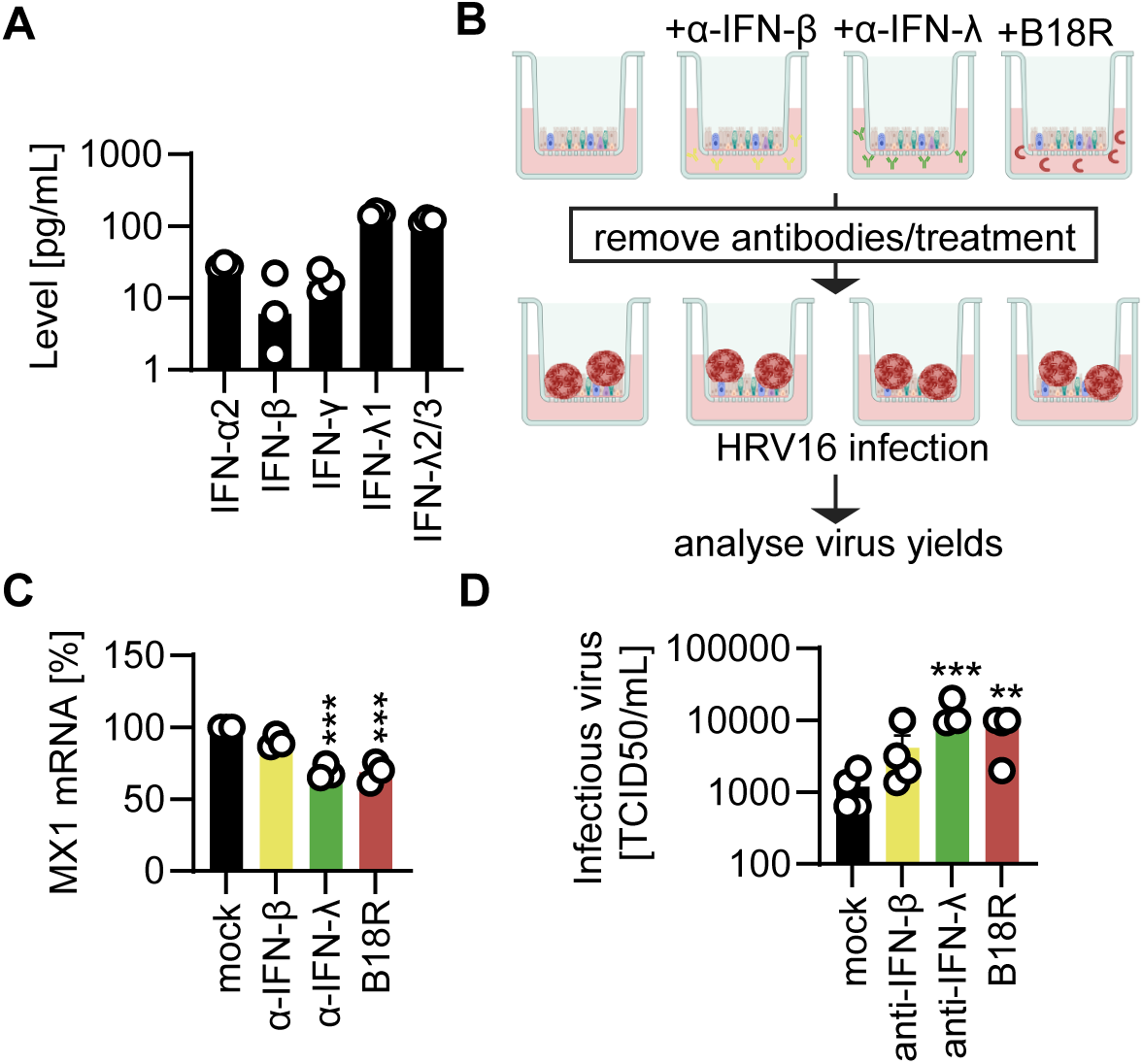
Lung epithelium tonic interferon serves as a major barrier against respiratory virus invasion. (**A**) Absolute levels of various IFNs, as indicated and detected via multiplex ELISA, in the basal supernatant of uninfected HBE models. Bars represent the mean of n=3±SEM (independent experiments). (**B**) Schematic representation of the workflow of tonic interferon depletion using neutralising antibodies (α-IFN-β (1 µg/mL); α-IFN-λ1/2/3 cocktail (1 µg/mL each of α-IFN-λ1,2 and 3) or scavenger (B18R, 200 ng/mL). Cultures were treated twice with a 24 h interval between treatments, each followed by a medium change. After 2 days treatment the cultures were thoroughly washed, infected with HRV (MOI 0.01) and apical washes collected in the following days. (**C**) Quantification of the MX1 mRNA levels 48 h indicated post-treatment of the HBE models as assessed by qPCR. Normalised to housekeeping gene TBP. Bars represent the mean of n=3±SEM (biological triplicates). One-way ANOVA with Dunnett’s multiple comparisons test. ***p = 0.0002. (**D**) Infectious HRV16 yields in the apical supernatants 1 dpi from HRV16 (MOI 0.01) infected HBEs assessed by TCID50. Bars represent the mean of n=4±SEM (independent experiments).

Summarizing, these findings indicate that tonic IFN signalling contributes to antiviral defence in the human bronchial epithelium, protecting against HRV16.

## DISCUSSION

Understanding host-pathogen interactions at the innate immune interface is crucial to understand the molecular pathogenesis but may also inspire preventive and cure strategies. To assess the molecular details, physiologically relevant models are key. Primary ALI cultures of the human lung epithelial cells are considered to reflect the natural respiratory interface. Indeed, our data shows a clear morphological similarity to the native tissue (Fig. 2).

A major limitation of previous protocols was the limited availability of primary donor cells and short virus growth times ∼7days (34, 44, 45). Here we introduced a method that tackled both these issues. Our data shows that we can amplify and expand the primary lung epithelial cells, for up to 12 passages without losing their differentiation potential (Fig. 1 and S1A-B). Furthermore, we showed that our primary ALI models supported long-term infection studies for up to 35 days (Fig. 4D). This suggests that our system could also be applied to evaluate antiviral toxicity, to define the dosing required for viral clearance, and to follow cellular fate after the infection has been cleared.

Cytokine profiling revealed a strong upregulation of host antiviral defences during early replication phase of HRV16. Consistent with our findings, both clinical and organotypic studies have also reported elevated early proinflammatory cytokine release during HRV16 infection (46–48). Notably, HRV16 could be cleared from the cultures in the absence of adaptive immune cells, suggesting that innate immune activation in the respiratory epithelium is sufficient to defend against HRV16. In contrast, the HCoV-229E and HCoV-NL63 were able to antagonise or delay the induction of type I and III IFNs response similar to SARS-CoV-2 (46–49). It is tempting to speculate that the robust and early IFN response likely facilitates clearance of HRV16, whereas evasion of this response enables the longer infections of common cold coronaviruses (53).

While the overall architecture and cell composition of the HBE and HSE model were similar, PCA analyses revealed differences in gene expression profiles. Interestingly, the HBE models expressed overall higher levels of mucosal defence genes compared to HSE (Fig. 3E-G). This difference may be attributed to structural and anatomical distinctions, as the bronchial epithelium is pseudostratified columnar whereas the small airway epithelium is predominantly cuboidal (54). Previous studies using airway epithelial cells have also reported variation in response in terms of cytokine release towards treatment between the different anatomical sites of the airway (32, 55).

Our data further show that tonic IFNs, in addition to infection-induced IFNs, play an important role in the airway epithelium’s antiviral defences. Infectious virus yields were more than 10-fold higher upon depletion of tonic IFNs (Fig. 5). This suggests that tonic IFNs may pre-activate barriers in potentially vulnerable tissues at the viral entry site to be ready for defence. Whether the levels of IFN in our model correspond exactly to those locally in the human lung remains open. Data from lung organoids derived from primary patient tissue suggest detectable cytokine levels in human lungs (30). In mouse models, tonic IFN levels, including IFN-γ usually range in the low pg/mL levels, similar to our detected IFN levels in the human HBE model (56, 57). Of note, tonic IFN levels are more studied in another mucosal surface, the gut (58, 59). Here, there were also shown to contribute to innate defences and immune homeostasis (59) or even hematopoiesis (60).

Several studies have proposed that tonic IFNs are mainly due to microbiota, often detected via cGAS-STING DNA sensing pathways (58, 59, 61, 62). For example, stimulation of innate defences by commensal microbiota through lung stroma or via dendritic cells may contribute to tonic IFN levels (63, 64) and anti-viral defence (62). However, our models are not colonised with bacteria and they do not possess resident macrophages or dendritic cells but still exhibit antiviral tonic IFN activity, thus their source is likely to be intrinsic. It was shown that basal IFN-λ helps shape the mucosal epithelial barrier even in the absence of pathogens in human epithelial cell models, driven by cGAS–STING mediated mitochondrial DNA (65). Loss of USP22 in the respiratory epithelium enhances STING-dependent STAT1 activation, IFN-λ secretion, and ISG expression even in the absence of an infection, leading to antiviral protection (66). Recent work also suggested that disruption of post-Golgi trafficking activates tonic cGAS-STING signalling in the absence of pathogenic stimuli enhancing basal interferon signalling and conferring antiviral and antitumor protection (67). With the integration of single-cell and spatial transcriptomic and proteomic approaches, future studies may delineate the epithelial subsets responsible for tonic IFN production and map viral receptor distributions that influence disease severity using our model.

## MATERIALS AND METHODS

### Primary cells, cell lines and media

Normal human bronchial epithelial cells (NHBE, Lonza, CC-2540, CC-2540S), and small airway epithelial cells (SAEC, Lonza, CC-2547) were cultivated in PneumaCult-Ex Plus complete media (StemCell Technologies, 05040) for expansion in 2D culture conditions. H1-HeLa-cells (ATCC #CRL-1958) were grown in Dulbecco’s Modified Eagle Medium (DMEM, Gibco, 41965039) containing 10% heat inactivated fetal bovine serum (FBS, Gibco, A5256701), 2mM Gluta-MAX™ (Thermo, 35050061) and 6.5 µg/mL Gentamicin (PAN-Biotech, 15710-049). Huh-7 cells (kindly provided by Anna-Laura Kretz, Department for General and Visceral Surgery, Ulm University) were grown in DMEM supplemented with 100 U/mL penicillin, 100 µg/mL streptomycin (PAN-Biotech, P06-07100), 2 mM L-glutamine (PAN-Biotech, P04-80100) and 10 % heat-inactivated FBS. Caco-2 cells (kindly provided by Holger Barth, Institute of Experimental and Clinical Pharmacology, Toxicology and Pharmacology of Natural Products, Ulm University Medical Center) were grown in DMEM supplemented with 100 units/mL penicillin, 100 µg/mL streptomycin, 2 mM L-glutamine, 1 mM sodium pyruvate (Thermo, 11360039), 1x non-essential amino acids (Gibco, 11140050) and 20 % heat-inactivated FBS. LLC-MK-2 cells (kindly provided by Lia van der Hoek, Laboratory of Experimental Virology, Department of Medical Microbiology, Amsterdam UMC, Location AMC, University of Amsterdam) were cultured in MEM (Sigma-Aldrich, M4655) supplemented with 100 units/mL penicillin, 100 μg/mL streptomycin, 2 mM L-glutamine, 1x non-essential amino acids and 8% FBS.

### Ethics

Organoids and ALI cultures were derived from fully anonymised, commercially available human primary epithelial cells (NHBE, Lonza, CC-2540, CC-2540S; SAEC, Lonza, CC-2547). Human bronchial tissue used for immunofluorescent staining was obtained from one donor (male, 48 years old) undergoing elective pulmonary resection at the University Hospital Würzburg. Informed consent was obtained beforehand, and the study was approved by the institutional ethics committee on human research of the Julius-Maximilians-University Würzburg (vote 182/10)

### Organoid generation and culture reagents

Epithelial cells were thawed and centrifuged at 800 g for 3 minutes. After removing the supernatant, the pellet was resuspended in BME Type II (R&D Systems, 3533-010-02) and plated as 50 µL domes/well in a pre-warmed 24-well plate (Sarstedt, 60823644). The plate was incubated at 37°C for approx. 15 min to allow the polymerization of BME Type II. The domes were then overlaid with 500 µL of organoid growth medium supplemented with 10µM Y27632 Rock-Inhibitor, only added freshly on the day of seeding (StemCell Technologies, 72304) per well. The organoid growth media was developed in-house based on previous research (68). It consists of Advanced DMEM-F12 (Thermo, 12634010) with the following factors 1X R-spondin1-CM (produced in-house), 1X GlutaMax (Gibco, 35050-038), 10mM HEPES (Lonza, 17-737E), 1X B27 supplement (Thermo, 12587-010), 5mM Nicotinamide (Merck, N3376-100G), 1.25 mM N-acetylcysteine (Sigma, A9165-5G), 100 µg/mL Primocin (Invivogen, ant-pm-2), 100 ng/mL mNoggin (Peprotech, 250-38), 25ng/mL FGF 7 (Peprotech, AF-100-19), 100 ng/mL FGF 10 (Peprotech, 100-26), 500 nM A 83-01 (Tocris, 2939), 10µM Y27632 Rock-Inhibitor, only added freshly on the day of seeding, 500 nM SB202190 (Selleckchem, S1077). For passaging, 500 µL of TrypLE express (Invivogen, 12604013) were added to each dome. Dome structures were destroyed by pipetting up and down and transferred into a 15 mL reaction tube (Sarstedt, 62554502). Organoids were incubated with TrypLE for up to 15 minutes at 37°C in a water bath. Singularization was supported by mechanical forces, pipetting up and down at least 15 times. Reaction was stopped by adding resuspension media to reach a final volume of 10 mL and cell number was determined. Upon centrifugation at 800 g for 3 minutes the cell pellet was resuspended in BME Type II and plated as 50 µL domes in a 24 well plate containing approximately 75,000 to 100,000 cells/dome. After polymerization for 15 min at 37 °C 500 µL of medium supplemented with 10 µM Y27632 were added.

### ALI model generation

Collagen (Stemcell technologies, 4902) was diluted 1:50 in Dulbecco’s Phosphate Buffered Saline (DPBS, Gibco, 14190-094) and 100 µL were added to 6.5 mm Transwell with 0.4 µm pore polyester membrane inserts (Corning, 3470). The plates were incubated over-night with lid open to dry the collagen and UV sterilised for 30 min on the following day. The plates with inserts were directly used for experiments or stored at 4°C until use. The surface of the inserts was washed with PBS before seeding the cells. For cell differentiation on trans well inserts, organoids were treated with TrypLE as described above and 1,000,000 cells were resuspended in expansion media containing 10 µM Y27632. 50,000 cells from dissociated organoids (NHBE and SAEC) were seeded on the inserts with PneumaCult-Ex Plus Medium (Stemcell Technologies, 05040) and cultured under submerged condition (300 µL media on apical compartment and 700 µL media on the basal compartment) until they were confluent (3 - 4 days). Media from the apical compartment was removed and the models were cultured under air-liquid interface (ALI) to induce cell-differentiation with 600 µL of complete PneumaCult-ALI medium (Stemcell Technologies, 05001), on the basal compartment. Fresh media was replaced thrice a week for 20 days and washed apically with 100 µL warm PBS once a week until muco-ciliary phenotype was observed (within 30 days after seeding).

### Viral strains and propagation

hCoV-229E-GFP (GFP gene in place of ORF4a, kindly provided by Volker Thiel, Department of Infectious Diseases and Pathobiology, Vetsuisse Faculty, University of Bern) was propagated in Huh-7 cells, hCoV-NL63 (kindly provided by Lia van der Hoek) on LLC-MK2 cells and HRV16 (ATCC #VR-283) on H1-HeLa-cells. To this end, 80 % confluent cells were inoculated with a MOI of 0.1 in medium supplemented with 2 % FCS and incubated at 33°C. Cells were monitored daily a the light microscope until a strong cytopathic effect was visible (4 days for hCoV-229E-GFP, 5 days for hCoV-NL63 and 3 days for HRV16). Supernatant was harvested and clarified by centrifugation at 1300 rpm for 5 minutes to remove cellular debris and stored at −80°C until use.

### TCID50 assay

Infectious titers of hCoV-229E-GFP, hCoV-NL63 and HRV16 were determined in 96 well format on Huh7 cells, Caco-2 cells or H1-HeLa-cells, respectively. 25,000 cells (Huh7, Caco2) or 20,000 cells (H1-HeLa) were seeded in 96 well plates in medium containing 2 % FCS. The next day, cells were inoculated with a 10-fold serial dilution of the virus stock and incubated at 33°C. At 7 days post infection, CPE was evaluated by light microscopy and TCID50 was calculated according to the Reed-Muench method (69)

### Live-cell video microscopy

Dynabeads Protein G particles (Invitrogen, 1004D) were mixed into DPBS to a final concentration of 30 µg/mL and pipetted 50 µL onto the models after apically washed with warm DPBS. Short videos of the transport of the particles across the surface of the models were recorded using high speed video microscopy (Leica. DMIL S80 PH FLUO).

### Immunofluorescence staining

The ALI cultures were fixed using 4 % PFA (600 µL at the bottom and 100 µL at the apical compartment) overnight at 4 °C and replaced with PBS and stored at 4 °C until use. The insert were inverted and the membranes containing the epithelial layer were gently removed using a scalpel and were either used directly for whole mount staining or were paraffin-embedded to make 5 µm sections on glass slides. For immunostaining, the paraffin was removed by placing the slides at 60 °C overnight, followed by washes with xylene (2x 10 min), 96% ethanol (2x 1 min), 70% ethanol (1 min), 50% ethanol (1 min) and distilled water (1 min). The sections were unmasked to expose the epitopes, post-embedding. The slides were immersed in sodium-citrate buffer (pH 6) or Tris-EDTA buffer (pH 9), depending on the antibody as recommended by the manufacturer, or optimised for each antibody and placed in a steam chamber for 10 min. The slides were retrieved and placed in deionised water for 3 min to cool before immunostaining. The FFPE sections or the whole epithelium for whole mount staining were permeabilised and blocked with 5% BSA in 0.5% TritonX-100 (Sigma-Aldrich, T8787) in PBS for 1 h to reduce the non-specific binding of the antibodies. The primary antibodies were diluted in the same blocking buffer and pipetted over the sections, and incubated overnight at 4 °C in a humidified chamber. The slides were washed three times in wash buffer (PBS containing 0.05% Tween-20 (Sigma-Aldrich, P9416)), followed by incubation with secondary antibodies and DAPI (Invitrogen, SBA010020) for 1 h at RT. The slides were rewashed three times, mounted, and covered with coverslips to prevent drying. The primary antibodies used were CK5 (1:200; Agilent, M7010), Acetylated tubulin (1:1000; Sigma-Aldrich, T8328), MUC5B (1:100; Sigma-Aldrich, HPA008246), MUC5AC (1:1000; Thermo, MA138223). The secondary antibodies used were Goat anti-Mouse IgG2b, Alexa Fluor 488 (1:400; Thermo, A-21141), Goat anti-Rabbit IgG (H+L), Donkey anti-Rabbit IgG, Alexa Fluor (1:400; Thermo, A-31572), Alexa Fluor 647 (1:400; Thermo, A-21245). Negative controls (omission of primary antibodies) were performed in each experiment to monitor non-specific binding. The samples were imaged using Zeiss LSM 710 confocal laser scanning microscope using the ZEN 2010 software, and Keyence BZ9000 E BIOREVO System and image analysis was conducted with ImageJ (42).

### RNA isolation and RT-qPCR

The RNA was isolated from the supernatants using QIAamp Viral RNA Mini kit (Qiagen, 52906) and cellular RNA was isolated using RNeasy Plus Mini Kit according to recommended protocol (Qiagen, 74136). To determine the HRV16 (Thermo, Vi99990017_po), 229E (Thermo, Vi06439671_s1) and NL63 (Thermo, Vi06439673_s1) levels, real-time quantitative PCR (RT-qPCR) was performed using TaqMan Fast Virus 1-Step Master Mix and on StepOnePlus Real-Time PCR System (96-well format, fast mode). qRT-PCR for cellular RNA was performed in one step using the Luna Universal Probe One-Step RT-qPCR (New England Biolabs Inc., E3006E) kit on a StepOnePlus Real-Time PCR System according to the manufacturer’s instructions. TaqMan probes for each individual gene [FOXJ1 (Hs00230964_m1), CK5 (Hs00361185_m1), MUC5AC (Hs01365616_m1), MUC5B (Hs00861595_m1), MX1 (Hs00895608_m1)] and housekeeping gene TATA-box binding protein gene (*TBP,* Hs00427620_m1), were acquired as premixed TaqMan Gene Expression Assays and added to the reaction. Expression level for each target gene was calculated by normalizing against TBP using the ΔCT method.

### NGS library preparation and RNA-seq analyses

RNA-seq libraries were prepared using the prime-seq protocol, a bulk RNA sequencing method that introduces barcodes during reverse transcription (RT) previously described (70). For each sample, 40 ng of total RNA were used as input for RT. After cDNA synthesis and amplification, sequencing libraries were constructed using the NEBNext Ultra II FS DNA Library Prep Kit for Illumina (NEB #E6177). Libraries were sequenced on an Illumina NovaSeq X (25B flow cell) with paired-end 150bp reads (Read 1: 150bp, Read 2: 150bp) and dual 8 bp indexes (Index 1: 8 bp, Index 2: 8 bp) targeting 5 million reads per sample. Raw reads were processed with the zUMIs pipeline (v2.9.7e) and aligned to the human reference genome GRCh38.p13 using STAR (v2.7.9a) with GENCODE v41 annotations (71, 72). The resulting count matrix was imported into R; read counts were filtered, normalized with DeSeq2, and variance-stabilized (vst) for downstream PCA and heatmap visualization. To estimate relative cell-type abundance from bulk RNA-seq, we used the CIBERSORTx web tool (73). A signature matrix was derived from single-cell RNA-seq data of healthy human airways and applied to impute cell-type fractions (14).

### Analyses of anti-viral response using LEGENDplex

The cytokines present in the basal compartment of ALI-cultures were quantified using the LEGENDplex Hu Anti-Virus Response Panel 1 (BioLegend, 741270) and its LEGENDplex Data Analysis Software. The assay and measurement by flow cytometry (FACS-Canto II) was performed according to the manufacturer’s instructions.

### Infections

The models were given fresh medium and washed apically with warm PBS (150 µL) and infected with 229E, NL63, and HRV16 (0.01 MOI). 50 µL of virus suspended in serum free media were added to the apical compartment and incubated at 37 °C. After 3-4 h, the airway epithelium was washed with 2x warm PBS and further cultured with or without respective inhibitors. Inhibitors were added at a concentration of 10 µM Remdesivir (Selleckchem, S8932), 20 µM Camostat mesylate (Merck, SML0057), and 3 µM Rupintrivir (Sigma-Aldrich, PZ0315) to respective wells. For short- (0, 2, 4 dpi), long-term (0, 2, 4, 7 and 16 dpi), and HRV16 clearance (0, 7, 14, 21 and 35 dpi) infection kinetics, the apical washes were collected and after RNA isolation, viral RNA was quantified by RT-qPCR.

### Transepithelial electrical resistance (TEER)

The barrier integrity of the HBE and HSE models was assessed with TEER measurements. The models were given apical wash with PBS and transferred to 600 µL PBS containing 24-well plates and the apical compartment was filled with 300 µL PBS. The measurements were made using Millicell ERS 3.0 Digital Volt-Ohm meter (Merck, MERS03000) in triplicates. The final TEER was expressed in Ω.cm^2^ by multiplying the values from Ohm meter x 0.33 (area of trans well insert).

### FITC-dextran epithelial barrier integrity assay

Fluorescein isothiocyanate (FITC) conjugated dextran (4 kDa; 46944-100MG-F, Sigma, Germany) flux across the apical to basal compartment was used as a read out for measuring paracellular transport and epithelial barrier integrity. The apical surface was washed with PBS to remove excess mucus, and 600 µL of fresh medium were added to the basal compartment. 100 μL of 0.25 mg/mL of FITC-dextran dissolved in Opti-MEM reduced serum media (Gibco, 31985047) sterile filtered using 0.4 µm filters were incubated on the apical compartment for 30 min protected from light in the incubator. 100 μL of the medium were collected from the basal compartment into a 96 well flat bottom plate in duplicates and absorbance at 490 nm was measured using Fluostar Omega. The mean absorbance measured in three independent experiments (N=3, in duplicates) was normalised to a cell-free trans well insert and the permeability of the human airway epithelial model to FITC-dextran was displayed as percentage values.

### Interferon depletion assays

The fully differentiated ALI cultures were washed apically with PBS and transferred onto fresh plates containing 600 µL medium with or without neutralising antibodies or inhibitor in biological triplicates. Two experimental time points were assessed, Day 0 to check for the basal ISG level and Day 1 to check for the viral RNA in each treatment groups post 24 h of HRV16 infection. Each time points had four treatment groups, treated either with Human IFN-β antibody (1 µg/mL; R&D, MAB814), or Human IFN-λ cocktail made by combining IFN-λ 1 antibody (1 µg/mL; R&D, MAB15981), IFN-λ 2 antibody (1 µg/mL; R&D, MAB1587), and IFN-λ 3 antibody (1 µg/mL; R&D, MAB5259) or B18R (200 ng/mL; Type I Interferon inhibitor) and untreated samples served as control. The models were treated for 48 h and 24 h with fresh media change in between treatments and apically washed to remove most of the tonic interferons. The Day 0 samples were immediately lysed after treatments (x2) and was subjected to RNA isolation and parallelly the infection group were infected with 0.1 MOI HRV16 for 24 h followed by cellular RNA isolation and apical supernatant collection for TCID50.

## Data availability

RNAseq data was deposited to Gene Expression Omnibus (NCBI), GSE312700. To review https://www.ncbi.nlm.nih.gov/geo/query/acc.cgi?acc=GSE312700 and enter token yrireewqhbyfxeb.

## Declaration of interests

The authors declare no conflict of interest.

## ACKNOWLEDGMENTS

We are grateful to Jana-Romana Fischer, Birgit Ott, Ralf Köhntop, Markus Jaritz (IMP Vienna) and the Vienna Biocenter Next Generation Sequencing Facility for excellent technical support. K.M.J.S. acknowledges funding by the German Federal Ministry of Education and Research (BMBF; IMMUNOMOD-01KI2014), the German Research Foundation (DFG; CRC1279, SP 1600/7-1, SP 1600/9-1) and the Baden-Wuerttemberg Stiftung (AutophagyBoost). R.S. was supported by a Baustein grant from the Medical faculty from the University of Ulm (L.SBN.0234). Work in the Gaidt lab was supported by Boehringer Ingelheim and the ERC (StG Guardians 101117146).

